# Meta-population structure and the evolutionary transition to multicellularity

**DOI:** 10.1101/407163

**Authors:** Caroline J. Rose, Katrin Hammerschmidt, Yuiry Pichugin, Paul B Rainey

## Abstract

The evolutionary transition to multicellularity has occurred on numerous occasions, but transitions to complex life forms are rare. While the reasons are unclear, relevant factors include the intensity of within-versus between-group selection that are likely to have shaped the course of life cycle evolution. A highly structured environment eliminates the possibility of mixing between evolving lineages, thus ensuring strong competition between groups. Less structure intensifies competition within groups, decreasing opportunity for group-level evolution. Here, using populations of the bacterium *Pseudomonas fluorescens,* we report the results of experiments that explore the effect of lineage mixing on the evolution of nascent multicellular groups. Groups were propagated under regimes requiring reproduction via a life cycle replete with developmental and dispersal (propagule) phases, but in one treatment lineages never mixed, whereas in a second treatment, cells from different lineages experienced intense competition during the dispersal phase. The latter treatment favoured traits promoting cell growth at the expense of traits underlying group fitness – a finding that is supported by results from a mathematical model. Together our results show that the transition to multicellularity benefits from ecological conditions that maintain discreteness not just of the group (soma) phase, but also of the dispersal (germline) phase.

## Introduction

Multicellular life evolved on independent occasions from single celled ancestral types. Explanations are numerous, ranging from those that emphasise the centrality of cooperation (Queller and Strassmann 2009; Bourke 2011; West et al. 2015), to perspectives that give prominence to specific mechanisms (Boraas et al. 1998; van Gestel and Tarnita 2017; Herron et al. 2019), through those who see vital ingredients residing in ecological factors that underpin emergence of Darwinian properties (Griesemer 2001; Rainey 2007; Godfrey-Smith 2009; Rainey and Kerr 2010; Libby and Rainey 2013a; De Monte and Rainey 2014; Rainey and De Monte 2014; Black et al. 2020).

Evidence for a seminal role for ecology comes from an on-going experiment that took inspiration from ponds studded with reeds and colonised initially with a planktonic-dwelling aerobic microbe. Growth of the microbe depletes oxygen, but the essential resource is available at the air-liquid interface. Growth at the meniscus requires production of adhesive glues (Spiers et al. 2002, 2003; Lind et al. 2017) that allows formation of mats comprised of sticky cells (simple undifferentiated collectives), but for mats to remain at the surface attachment to a reed is required. Attachment of genetically distinct mats to different reeds ensures variation among mats. From time to time a mat detaches from a reed and sinks. Death provides opportunity for an extant mat to export its success to a fresh reed (as long as some means of dispersal is possible). As a consequence of patchily distributed resources and a means for mats to disperse among reeds, a Darwinian-like process stands to unfold at the level of mats (Rainey and Kerr 2010; Rainey et al. 2017; Black et al. 2020).

The experimental evolution analogy uses the bacterium *Pseudomonas fluorescens* and glass microcosms as a proxy for reeds (Hammerschmidt et al. 2014). Growth of non-sticky (smooth (SM)) planktonic cells depletes oxygen from the broth phase, establishing conditions that favour the evolution of mat-forming (wrinkly spreader (WS)) cells. Formation of mats establishes conditions that favour the further evolution of non-sticky cells within the mat (Rainey and Rainey 2003). Continuing time-lagged frequency dependent interactions between SM and WS types (Rainey and Travisano 1998) generates a simple life cycle (Libby and Rainey 2013b) that becomes the focus of selection (Figure 1a).

**Figure 1.**
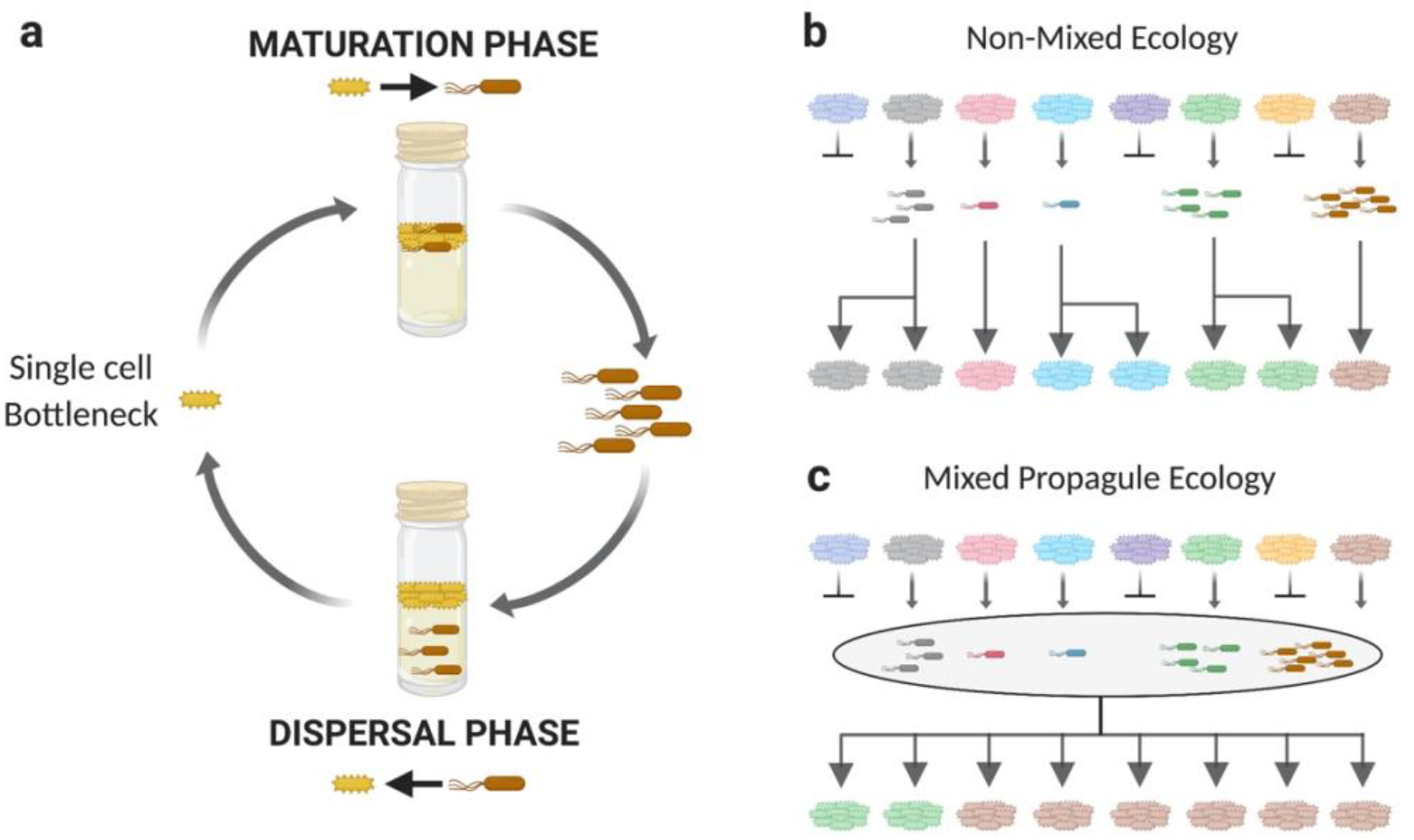
Experimental Regimes. (a) A single ‘mat’ generation consists of a life cycle of two phases. The Maturation Phase is seeded with a single WS cell. SM cells arise within the mat and are harvested after six days of maturation by plating and collection of all SM colonies on agar plates. The SM propagule cells are transferred to a new microcosm to begin a three-day Dispersal Phase, during which WS mat-forming cells arise. At the end of the Dispersal Phase, cells from microcosms are plated once more, and a single WS colony (representative of the most common colony morphology type) is picked to seed the next generation. Mat extinctions occur if there are no SM cells after six days of the Maturation Phase, no WS cells after three days in the Dispersal Phase, or if the mat collapses during the Maturation Phase. (b) and (c) Schematic depiction of a population of eight genetically distinct groups (indicated by different colours) proceeding through one life cycle within their respective non-mixed (b) and mixed (c) ecologies.

Because the cycle is initially dependent upon spontaneous mutation, it is prone to failure (but lines can also fail through production of fragile mats). Lineages that fail are removed, thus allowing extant types to export their success to new microcosms in precisely the same way as a mat that falls from a reed provides opportunity for competing mats to export their reproductive success. The non-sticky motile cells act as dispersing agents analogous to a germ-line. The mat itself serves both an ecological role by ensuring access to oxygen, while also producing seeds for the next generation of mats. In this regard, the mat, in the absence of non-sticky dispersing cells is analogous to soma (and an evolutionary dead-end).

After ten life cycle generations, mats propagated under the two-phase life cycle regime evolved – in one lineage – a simple genetic switch that reliably transitioned successive life cycle phases, but more striking was the overall impact of the longer timescale (the nine-day time required for doubling of mats) on the shorter timescale (the hourly doubling of cells). As shown by Hammerschmidt et al. (2014), selection over the long timescale caused the fitness of mats to increase (as determined by the relative ability of mats to give rise to offspring mats), while fitness of the individual cells comprising mats declined (when measured relative to ancestral types). This can be understood in terms of selection over the longer timescale trumping the effects of individual cell selection: over the long-term, successful cells are those whose fitness aligns with the longer timescale defined by the longevity of the nascent multicellular organism (Bourrat, 2015; Black et al. 2020). Such an alignment of reproductive fates during the transition from cells to multicellular organisms has been referred to as “fitness decoupling” (Michod and Roze 1999) – a term that captures the sense that when selection comes to act over the longer timescale, fitness of the lower level particles “decouples” from that of the higher level collective.

Included in the experiment was a second treatment where mats evolved with a life cycle involving just a single phase: mats gave rise to mat-offspring via a single sticky mat-forming cell. After ten life cycle generations mat fitness improved, but there was no evidence of fitness decoupling: enhanced fitness of mats was readily explained by enhanced fitness of individual cells (Hammerschmidt et al. 2014; Rose et al. 2020).

This result drew particular attention to the significance of the two-phase life cycle. For the evolution of multicellular life – given appropriate ecological circumstances – such a life cycle delivers in a single step a second time scale (Black et al. 2020) over which selection might act (replete with birth-death events), a developmental programme that stands to become the focus of selection, a reproductive division of labour, and even the seeds of a distinction between soma and germ.

One might reasonably ask whether, if such life cycles can arise with such seeming ease, why multicellularity hasn’t arisen more often. One possibility is that ecological conditions are more restrictive than indicated by the reed / pond analogy. In fact, in the regime implemented by Hammerschmidt et al. (2014), lineages never mixed: mats were founded by single cells with discreetness maintained by virtue of boundaries afforded by the microcosms, similarly, dispersing cells from each mat were maintained as separate lineages. In the reed / pond analogy, dispersing cells arising from different mats are released into the planktonic phase and are thus expected to compete with a diverse range of dispersing genotypes. This subtle distinction is likely important.

Here we explore the impact of population structure on the emergence of individuality. The life cycle from our previously published results (“Non-Mixed Ecology” treatment; Figure 1b) is contrasted with an identical two-phase life cycle that incorporates competition (mixing) during the dispersal phase. This environmental manipulation, which is here termed the “Mixed Propagule Ecology” treatment (Figure 1c), was performed simultaneously with the earlier study. The results show that competition effected during the dispersal phase of a two-stage life cycle leads selection to favour traits that promote cell growth at the expense of traits underlying group fitness. This conflict is due to a tradeoff between traits underlying the fitness of groups and their constituent cells, and is supported by findings derived from a mathematical model. While the existence of a germ line can bring about the decoupling of fitness required to achieve a higher level of individuality, intense competition between propagule cells skews selection towards traits that enhance the competitive ability of cells, rather than towards traits that enhance group function, to which the life cycle is integral.

## Methods

### Experimental regime

The Non-Mixed Ecology treatment has been previously published in a study that compared its effect relative to a life cycle without reproductive specialisation (Hammerschmidt et al. 2014). Here the effect of meta-population structure on the evolutionary transition to multicellularity is addressed. Groups of cells (‘microcosms’) in both the Non-Mixed and Mixed Propagule treatments of the present study experience identical two-phase life cycles driven by frequency-dependent selection. More specifically, each of the Non-Mixed and Mixed Propagule meta-population ecologies comprised 15 replicates of eight competing groups that were founded with *P. fluorescens* strain SBW25 (Silby et al. 2009), and propagated through ten generations of evolution (one generation equated to one WS-SM-WS life cycle (Hammerschmidt et al. 2014).

Maturation Phase (Figure 1): Each group was founded by a single WS colony. Microcosms were incubated under static conditions for six days, after which they were checked for the presence of an intact mat at the air-liquid interface. If the mat was not intact, that line was deemed extinct.

Dispersal Phase (Figure 1), Non-Mixed Ecology: All microcosms with viable mats were homogenised by vortexing and then individually diluted and plated on solid media. Agar plates were subsequently screened for SM colonies. Lines without SM colonies were deemed extinct. To ensure that only SM cells, and no WS cells, were transferred to the Dispersal Phase, all SM colonies were individually transferred to 200 μl liquid medium and incubated for 24 h under static conditions. Thereafter they were pooled and used to inoculate Dispersal Phase microcosms. Each mixture of SM cells arising from within each individually plated microcosm was used to inoculate one or more daughter microcosms in the Dispersal Phase. When a microcosm was deemed extinct at the end of the Maturation Phase, it was replaced by a pool of SM (dispersing) cells from another microcosm randomly chosen from the same population of eight (Figure 1b).

Dispersal Phase (Figure 1), Mixed Propagule Ecology: All microcosms with viable mats were homogenised by vortexing and then pooled prior to diluting and plating on solid media. Agar plates were subsequently screened for SM colonies. To ensure that only SM cells, and no WS cells, were transferred to the Dispersal Phase, all SM colonies were individually transferred to 200 μl liquid medium and incubated for 24 h under static conditions. Thereafter they were pooled and used to inoculate Dispersal Phase microcosms. Because all microcosms with viable mats were pooled prior to plating, only one SM mixture was generated, and this mixture was used to inoculate all eight microcosms entering the Dispersal Phase (Figure 1c).

After three days of incubation under static conditions (during which new WS mats emerged), all microcosms in both treatments were individually plated on solid agar. The most dominant WS morphotype on each agar plate was selected to inoculate the next generation of the life cycle. If there were no WS colonies on the plate, the microcosm was deemed extinct. Figures 1b and 1c contrast the death-birth process of group competition in the Non-Mixed Ecology, with the physical mixing mode of competition in the Mixed Propagule Ecology.

### Fitness assay

Cell-level and group-level fitness were assayed after ten life cycle generations: 15 representative clones (one per replicate population) were generated from each of the evolved treatments, in addition to 15 ancestral WS lines (each independently isolated from the earliest mats to emerge from the ancestral SM strain SBW25) (described in detail in (Hammerschmidt et al. 2014)). For each genotype, three replicate competition assays were performed in populations of eight microcosms over the timescale of one full life cycle (Figure 1a) against a neutrally marked ancestral competitor (Zhang and Rainey 2007). In order to include the effects of both cell fitness and group fitness on the outcome of competition, all fitness assays were performed in the Mixed Propagule Ecology (Figure 1c). To simulate a meta-population structure with eight competing groups, four started with the marked reference strain and four started with the focal clone, the “SM mixture” used to inoculate the Dispersal Phase contained an equal volume of the marked reference strain for all focal strains. This ensured that the reference strain performed equally for each competition during the Maturation Phase. The single-celled bottleneck ensured that non-chimeric mat offspring could be counted at the end of the life cycle. Our proxy for group-level fitness is the proportion of ‘offspring’ mats produced at the end of one life cycle by the focal genotype relative to the marked reference strain, and cell-level fitness the total number of cells in the mat at the end of the Maturation Phase.

### Life cycle parameters

Density of WS and SM cells, and Proportion of SM cells were also assayed at the end of the Maturation Phase. The growth rate of SM cells was determined from three biological replicate SM colonies per line (for details on how the SM were obtained, see (Hammerschmidt et al. 2014)) in 96-well microtitre plates shaken at 28°C, and absorbance (OD600) measured in a microplate reader (BioTek) for 24 h. The experiment was repeated three times and the maximum growth rate (Vmax) was calculated from the maximum slope of absorbance over time. The transition rate between WS and SM cells, i.e., the level of SM occurrence in the Maturation Phase, and WS occurrence in the Dispersal Phase, was determined in a separate experiment, where static microcosms were individually inoculated with single colonies of the representative WS types. The Maturation Phase was extended from 6 to 12 days, and the Dispersal Phase from 3 to 6 days. At day six of the Maturation Phase, SM cells were collected to inoculate microcosms for the Dispersal Phase. Each day, three replicate microcosms per line were destructively harvested. The number of microcosms with SM cells was noted, and the number of SM and WS colony forming units determined.

### Statistical analysis

For detecting differences in group-level fitness and transition rate between cells of the evolved and ancestral lines, generalized linear models (error structure: binomial; link function: logit) with the explanatory variables Ecology, and representative clone (nested within Ecology) were calculated. Analyses of variance (ANOVA) were used to test for differences in cell-level fitness, density of WS cells, and density, proportion, and growth rate of SM cells between the evolved and ancestral lines. Explanatory variables were Ecology, and representative clone (nested within Ecology). Posthoc tests revealed differences between the evolved and ancestral lines. Relationships between the traits and cell and group-level fitness were tested using the mean per representative type accounting for regime. Pearson correlations and regressions were performed. The sample size was chosen to maximise statistical power and ensure sufficient replication. Assumptions of the tests, i.e., normality and equal distribution of variances, were visually evaluated. All tests were two-tailed. Effects were considered significant at the level of *P* = 0.05. All statistical analyses were performed with JMP 9. Figures were produced with GraphPad Prism 5.0, Adobe Illustrator CC 17.0.0, Inkscape 0.92.3 and Biorender.com.

### Model of selection regimes

The model simulates the evolutionary dynamics of metapopulations composed of *M* = 8 groups. Each group contained one or more lineages, which are the primary agents of selection. Each lineage in the metapopulation is characterized by three parameters: cell growth rate *ω*, transition probability *p*, and the number of cells in lineage *n*(*t*). At the beginning of each simulation, each group in a metapopulation was seeded with a unique lineage. The growth rate and transition probability of each lineage were sampled from a bivariate normal distribution with means 〈*ω*〉 = 1 and 〈*p*〉 = 10^-6^, variances σ_ω_ = 0.5 and *σ_p_* = 5 · 10^-7^, and correlation coefficient (between growth rate and transition probability) *ρ* = −0.5. The initial population of each lineage was set to a single cell. The dynamics of growth during Maturation and Dispersal Phases were simulated identically. Lineages grew exponentially according to their growth rates *ω_i_* until their combined size Σ_*i*_*π_i_* reached the carrying capacity of the group *N* = 10^6^ cells. Since each cell division in lineage *i* can result in a switch between phenotypes with probability *p_i_*, the number of phenotype switches during growth was sampled from Poisson distribution with rate parameter *p_i_π_i_*. The size of a lineage at which a phenotype transition event occurred 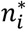 was sampled from a uniform distribution between one and *π_i_*. The moment at which this event occurred was calculated as 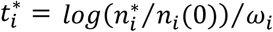. Each phenotype switch event resulted in the emergence of a new lineage of another phenotype, with growth rate and transition probabilities equal to those in the maternal lineage. The newly emerged lineages also grew exponentially and were sampled only at the end of the growth phase. At the end of each Dispersal Phase of the life cycle, a single novel lineage phenotype was sampled with probability proportional to its representation within its group. At the end of each Maturation Phase, all novel phenotype lineages were sampled in numbers proportional to their sizes. Each sample seeded one group at the beginning of the next growth phase. However, groups in which no phenotype switch events occurred did not contribute any samples at the end of the growth phase. These groups were deemed extinct and were reseeded by another random sample from the metapopulation. Seeding after the Maturation Phase differed in the Mixed Propagule Ecology: all samples were pooled together and the resulting mixture of lineages seeded all groups for the next Dispersal Phase. For both ecologies, simulations lasted for 20 full cycles and 600 independent realizations were performed. The average growth rate and transition probabilities across all groups were recorded for each simulation run.

## Results and Discussion

We begin with a brief description of the contrasting Non-Mixed Ecology and Mixed Propagule Ecology life-cycle regimes. Each generation began with a single WS cell, which through cell-level replication formed a mat at the air-liquid interface (Maturation Phase in Figure 1a). For a mat to reproduce it was required to be both viable and fecund, i.e., it had to produce SM propagule cells. In both ecological scenarios, competition between groups arose from a death-birth process: following an extinction event, a group was randomly replaced by a surviving competitor group. Extinction/replacement of groups occurred with high frequency (usually due to the lack of SM production (Hammerschmidt et al. 2014)), and therefore imposed potent between-group selection.

The two experimental treatments differed solely in manipulation of the Dispersal Phase of the life cycle (Figure 1b and 1c). In the Non-Mixed Ecology, SM cells were harvested separately from each surviving group at the end of the Maturation Phase. By contrast, in the Mixed Propagule Ecology, SM cells were harvested and *pooled* from all groups that survived the Maturation Phase. The pooled mixture was then used to seed all eight groups in the Dispersal Phase. In both ecologies, SM propagule cells competed within individual microcosms during the Dispersal Phase to produce WS types, and ultimately for mat formation. At the end of the Dispersal Phase, one colony of the most numerous WS morphology occurring in each microcosm was transferred to a fresh microcosm to begin the Maturation Phase of the next mat generation. Importantly, this step was performed for both treatments to ensure that all mats at the start of each new generation were seeded from a single cell.

### Changes in group and cell fitness

After ten group generations, changes in both cell and group level fitness were compared with a set of ancestral lines. Given the wide range of mutational pathways for evolution of WS from SM (McDonald et al. 2009; Lind et al. 2015; Lind et al. 2019), a range of ancestral WS lines was generated for comparison with the evolved lines. Each ‘ancestral’ line was a WS genotype isolated independently from the first mats emerging from the common SM ancestor (see Methods and (Hammerschmidt et al. 2014)) – this enabled a comparison of the distributions of fitness and other parameters of evolved and ancestral lines.

Fitness of all evolved and ancestral lines was estimated by competition in populations of eight microcosms with a common neutrally marked reference SM genotype (Zhang and Rainey 2007) over the timescale of one generation of the mat life cycle (Figure 1c). The single-celled bottleneck ensured that non-chimeric mat offspring could be counted at the end of the life cycle. Group fitness was the proportion of offspring mats produced (in a population of eight mats) by the focal line relative to the marked competitor, while cell fitness was the mean total number of cells present in the microcosms at the end of the Maturation Phase.

Fitness of derived lineages in the Non-Mixed Ecology significantly increased (ability to leave group offspring) relative to the ancestral types (*χ*^2^=32.660, d.f.=1, *P*<0.0001; Figure 2a), whereas cell fitness (number of cells present immediately prior to dispersal) decreased (*F*_1_ = 10.612, *P* = 0.002; Figure 2b). In contrast, under the Mixed Propagule Ecology, group fitness did not change (*χ*^2^=3.137, d.f.=1, *P*=0.077; Figure 2a), whereas cell fitness increased (*F*_1_ = 56.214, *P* < 0.0001; Figure 2b).

**Figure 2.**
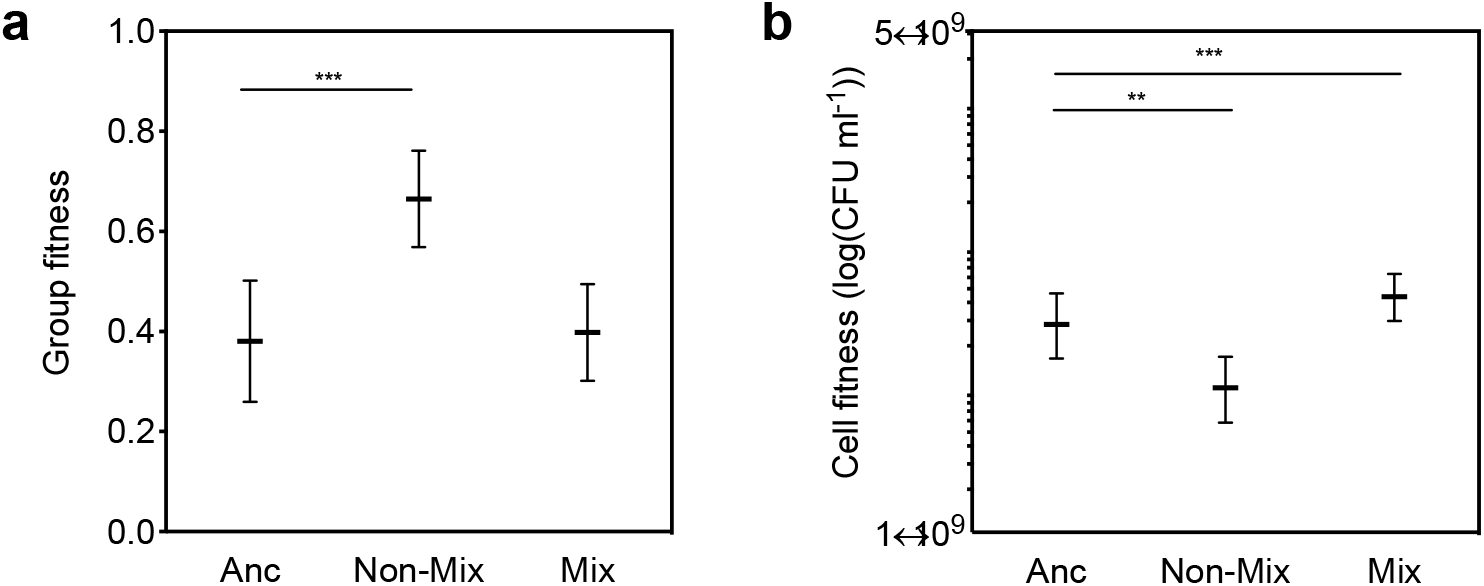
Changes in group (a) and cell (b) fitness in the Non-Mixed Ecology (Non-Mix) and Mixed Propagule Ecology (Mix) regimes compared to ancestral populations (Anc). Group fitness is the proportion of derived offspring mats after one life cycle relative to a genetically marked reference genotype. Error bars are s.e.m., based on n = 14 (Non-Mix) and n = 15 (Anc, Mix). ** denotes significance at the level of *P* = 0.001 – 0.01, and *** at the level of *P* <0.001.

At first glance, this is a surprising result. The group (WS mat) phase was identical in both treatments (each group was founded from a single WS cell). The only difference was the extent of competition among propagule cells. Under the Mixed Propagule Ecology there was no evidence of fitness decoupling as previously reported for the Non-Mixed Ecology (Hammerschmidt et al. 2014): there was no change in group fitness (relative to the ancestral type), but fitness of cells increased. Competition among single cells that comprise the propagule phase thus markedly affected the evolutionary fate of the evolving lineages. Such an effect draws attention to the fact that the evolving entities are defined by a life cycle comprised of both soma- and germ-like phases, and not simply by the group (WS) state. In the next sections we unravel the underlying causes, beginning with analysis of the ancestral state.

### Tradeoff between group and cell fitness

Figure 3a illustrates a negative relationship between cell and group fitness in the ancestral lines (*χ*^2^=4.246, d.f.=1, *P*=0.0393). It also shows evidence of a bimodal distribution of group fitness, indicative of a tradeoff between traits underpinning cell and group fitness.

**Figure 3.**
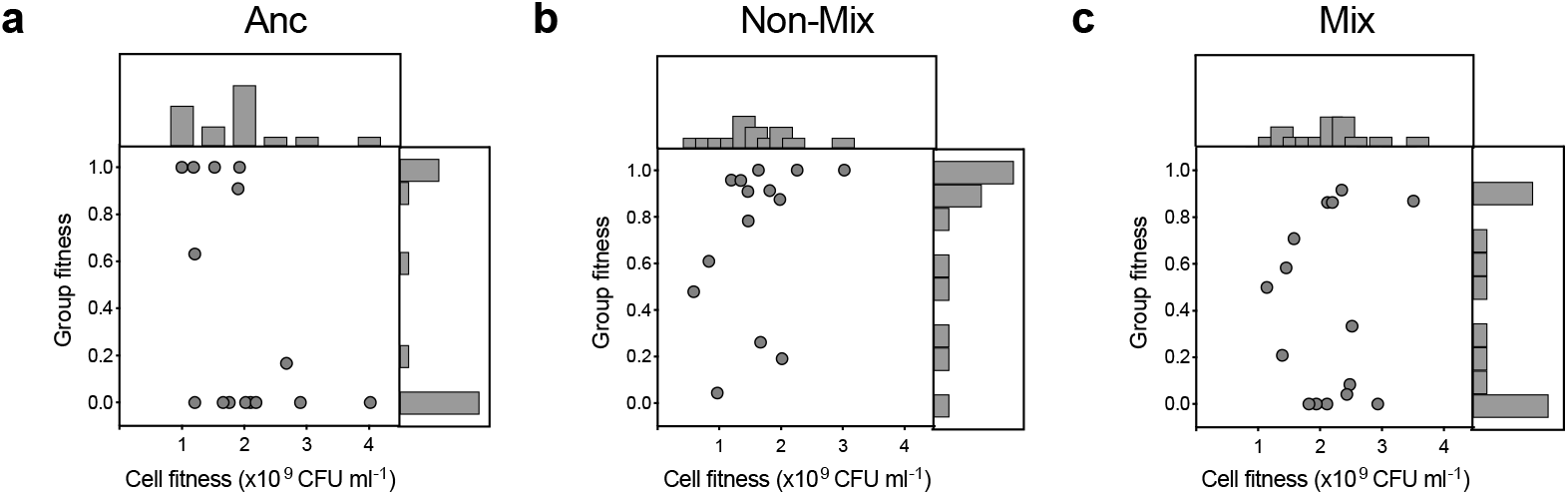
Relationship between cell and group fitness in the Non-Mixed (b) and Mixed (c) Propagule Ecologies compared to ancestral (a) populations. Group fitness is the proportion of derived offspring mats relative to a genetically marked reference genotype. Each dot represents the mean of eight lines per replicate population, assessed in three independent competition assays.

Ten generations of selection in the Non-Mixed Ecology shifted the distribution towards the ‘high group fitness / low cell fitness’ corner of the graph (Figure 3b), indicating that group-level selection was more potent than cell-level selection. Under the Mixed Propagule Ecology there was no corresponding change in the relationship between group and cell fitness in the derived lineages (Figure 3c).

The contrasting responses are most readily understood in terms of differences in the intensity of within-versus between-lineage selection. In the Non-Mixed Ecology regime lineages never interact during the Dispersal Phase and thus, competition -wrought via the death and birth of groups – occurred almost exclusively between lineages. Under the Mixed Propagule Ecology regime, while the WS mats that initiate the Maturation Phase were discrete and did not mix, the propagules collected after six days and used to found the Dispersal Phase were a pooled mixture sampled from each of eight microcosms. Thus, during the Dispersal Phase within-microcosm competition is intense, and appears to have overwhelmed between-lineage competition.

A further factor impacting the Mix-Propagule Ecology, and especially the opportunity for between-lineage selection, was reduced between-lineage variation. This was not directly measured, but was inferred from the identical visual appearance of WS mats in microcosms at end of the Dispersal Phase under the Mixed, but not Non-Mixed, propagule ecologies.

The causes of the reduced between-lineage variation are easily understood and worthy of consideration because they reflect a rarely considered downside of the standard trait group framework (Wilson 1975). Trait group models provide an explanation for the evolution of maintenance of behaviours that are costly to individuals, such as cooperation. Two genotypes are typically assumed: co-operators and defectors. The trait group model assumes that these types are randomly assembled into groups. Within groups, defecting types out-compete co-operators, but groups comprised of co-operators are more productive than groups dominated by defectors. Provided there is periodic mixing of the contents of all groups into a single global pool, followed by random assortment into new groups, then cooperation can be maintained. In essence group selection rewards those groups producing the largest numbers of individuals.

In the Mixed Propagule Ecology there is also a significant reward to WS mats that maximise production of SM propagule cells. But it comes with a cost to the efficacy of selection between groups. Consider a single WS type in one of eight microcosms that acquires an early mutation to SM and which therefore yields a vast excess of SM relative to each of the other seven WS mats. This successful SM type is thus overrepresented in the pool of SM propagules, which means that each of the eight microcosms that start the Dispersal Phase also contain an excess of this single genotype. Being more numerous, cells of this lineage are likely to be the source of the next WS-causing mutation. Furthermore, mutational biases arising from features of the genotype-to-phenotype map underpinning the transition between SM and WS types (McDonald et al 2009, Lind et al 2015, 2019), means that not only is it likely that the next WS type in each of the eight microcosms arises from the same SM lineage, but also arises via the exact same mutation, or at least a mutation in the same gene. The overall effect is to eliminate variation between groups, thus essentially eliminating the possibility of between-lineage selection.

### Changes in life cycle parameters

To identify traits contributing to differences in fitness between lineages subject to the non-mixed and mixed ecologies, we measured properties of WS mat and SM propagule cells expected to determine successful multicellular life cycles. After ten life cycle generations under the Non-Mixed Ecology regime, there was no change in the density, proportion, or growth rate of SM cells (measured at the end of the Maturation Phase) (density: *F*_1_ = 1.278, *P* = 0.2663; proportion: *F*_1_ = 2.702, *P* = 0.1095; growth rate: *F*_1_ = 2.116, *P* = 0.1522), however, the density of WS cells decreased (*F*_1_ = 8.036, *P* = 0.0065), while the rate of transition between WS and SM cells dramatically increased (χ^2^=114.198, d.f.=1, *P*<0.0001).

Evolution under the Mixed Propagule Ecology regime led to a reduction in the density and proportion of SM cells (density: *F*_1_ = 56.214, *P* < 0.0001; proportion: *F*_1_ = 102.217, *P* < 0.0001; growth rate: *F*_1_ = 2.664, *P* = 0.1103; Figure 4a,b,c), but an increase in the density of WS cells (*F*_1_ = 9.904, *P* = 0.0027; Figure 4d. Additionally, there was an increase in the rate of transition between WS and SM cells, but this did not approach the magnitude of the effect observed for the Non-Mixed Ecology (χ^2^=12.459, d.f.=1, *P*=0.0004; Figure 4e).

**Figure 4.**
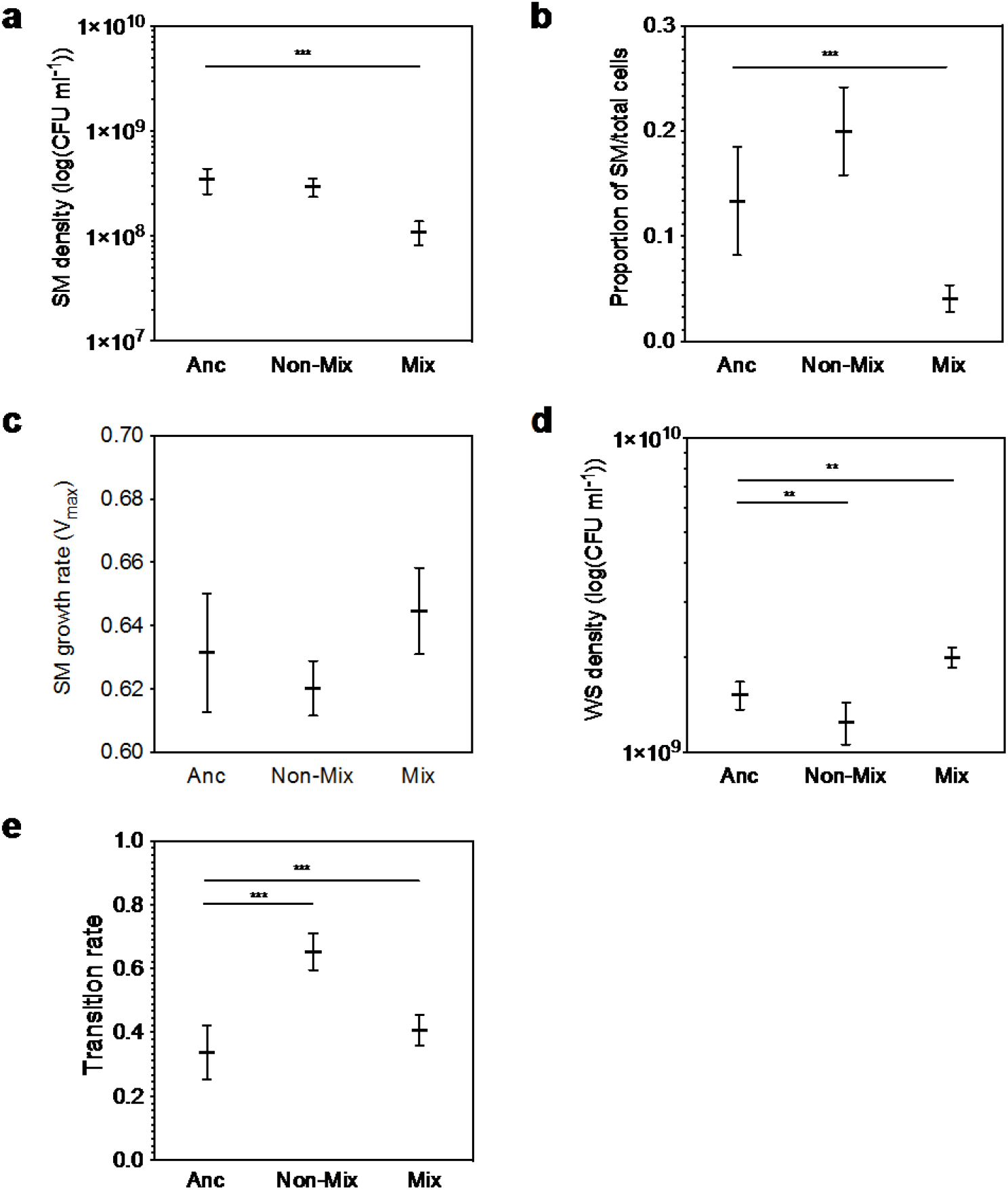
Changes in life cycle traits in the Non-Mixed (Non-Mix) and Mixed Propagule (Mix) Ecologies compared to the ancestral populations (Anc): (a) SM density, (b) Proportion of SM, (c) SM growth rate, (d) WS density, (e) Transition rate. Error bars are s.e.m., based on n = 14 (Non-Mix) and n = 15 (Anc, Mix). ** denotes significance at the level of *P* = 0.001 – 0.01, and *** at the level of *P* <0.001.

Understanding the connection between these data and the effects of selection wrought by the two contrasting ecologies is complex. A starting point is to recognise that under both treatment regimens the primary determinant of success is ability of lineages to generate each phase of the life cycle and critically to transition between phases. Given the importance of capacity to transition between states, the dramatic response in the Non-Mixed Ecology is not surprising, however, it is surprising that this response was so reduced in the Mixed Propagule Ecology (Figure 4e).

As mentioned above, a key difference is the extent of competition between propagules. Under the Mixed Propagule Ecology, propagules arising from mats during the six-day maturation phase must compete directly with propagules derived from other lineages during the dispersal phase. Given that the dispersal phase ends with sampling of a single WS colony (of the most common type) from each microcosm, representation in the next generation is thus determined solely by the number of WS cells at end of the dispersal phase. While this could in principle be achieved by increases in the growth rate or density of SM cells, the selective response was specific to the density of WS cells. In contrast, WS cells arising in the Dispersal Phase of the Non-Mixed Ecology need only outcompete any alternative WS cell types that may (or may not) arise within the same group. Thus, overall, mixing of propagules shifts the emphasis of selection from a developmental programme (capacity to transition through phases of the life cycle), toward density of WS cells (Fig. 4d). Moreover, the fact that under the Mixed Propagule Ecology, transition rate improved only marginally after 10 life cycle generations, whereas WS density significantly increased, points at a tradeoff between WS density – and by extension WS growth rate – and ability to transition through phases of the life cycle.

### Identification of traits linked to group and cell fitness

The fact that the WS-SM cell transition rate was the only measured parameter to increase in the Non-Mixed Ecology led to recognition that the WS-SM transition rate is associated with group fitness (Figures 5a – c). Indeed, these two factors are positively correlated in the ancestral lines (χ^2^=28.029, d.f.=1, *P*<0.0001; Figure 5a). During evolution in the Non-Mixed Ecology, the distribution shifted towards the ‘High Group Fitness/High Transition Rate’ corner of the spectrum with the two parameters still associated (χ^2^=13.657, d.f.=1, *P*=0.002; Figure 5b).

**Figure 5.**
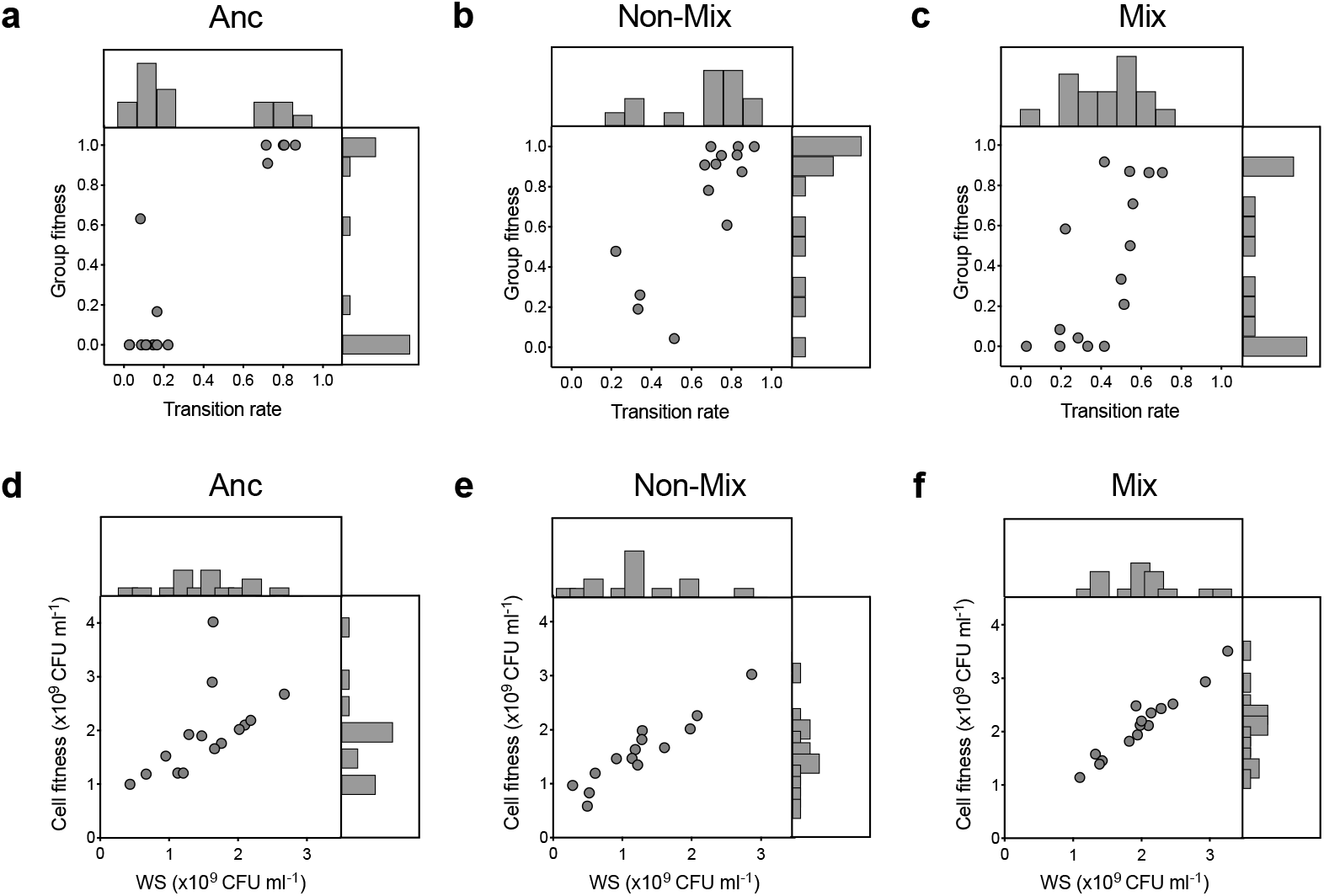
Relationship between life cycle traits and group and cell fitness. (a-c) Association of transition rate and group fitness in the ancestral populations (a), and in the Non-Mixed (b) and Mixed Propagule (c) Ecologies. (d-f) WS density is positively associated with cell fitness (total number of cells) in the ancestral populations (d), and in the Non-Mixed (e) and Mixed (f) Propagule Ecologies. Group fitness is the proportion of derived offspring mats after one lifecycle relative to a genetically marked reference genotype. Dots represent the mean of eight lines per replicate population, which were assessed in three independent competition assays.

Cell Fitness in the ancestral lines was strongly associated with the Density of WS cells (*F*_1_ = 6.673, *P* = 0.023; Figure 5d). The distribution of both parameters increased during the Mixed Propagule Ecology (*F*_1_ = 200.931, *P* < 0.0001; Figure 5f) and decreased during the Non-Mixed Ecology (*F*_1_ = 97.359, *P* < 0.0001; Figure 5e).

### Tradeoff between WS-SM cell transition rate and WS density

A negative relationship (tradeoff) exists between WS Density (which is linked to cell fitness) and WS-SM transition rate (which is linked to group fitness) in the ancestral population (r=-0.705, *P*=0.003, N=15; Figure 6a). The nature of the association explains both the negative relationship between the two levels of fitness observed above (Figure 3), and the opposing direction of selection in the two ecologies. While cells were required to survive an identical two-phase life cycle regardless of metapopulation structure, these two traits were driven in opposite directions under the two ecologies because of differences in the emphasis of cell and group level selection (Figures 6b, c).

**Figure 6.**
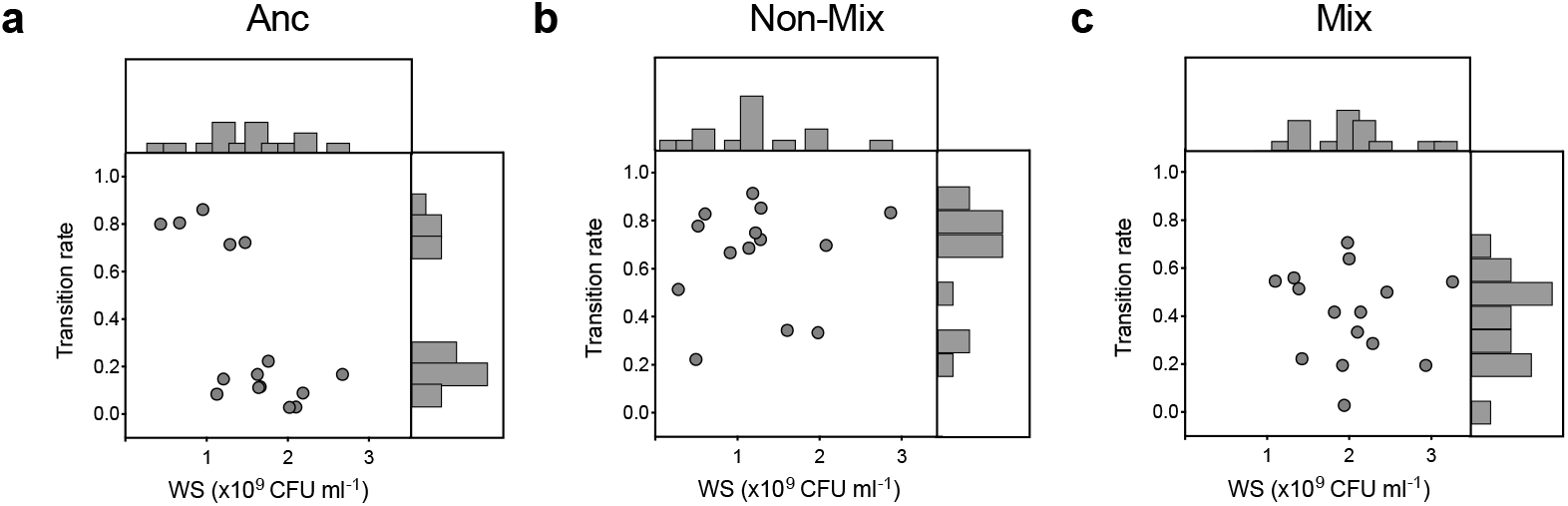
Relationship between WS density and transition rate in the ancestral populations (a), and in the Non-Mixed (b) and Mixed (c) Propagule Ecologies. Dots represent the mean of eight lines per replicate population, which were assessed in three independent assays.

### A simple model embracing cell- and group-level tradeoffs

To explore the extent to which the divergent evolutionary trajectories of groups evolving under the non-mixed and mixed regimes might be attributed to the experimentally recognized tradeoff between cell and group fitness, and more specifically density of WS cells (the cell-level trait) and transition rate (the group trait), a simple model of group-structured populations was developed (see Methods for details).

In the model, cells are characterized by two quantitative traits: growth rate and probability of transitioning between phenotypes. Independent lineages, with parameters drawn randomly from a bivariate normal distribution, found each group. The tradeoff between cell and group fitness observed in the ancestral bacterial population (Figure 3a) was implemented using a trait distribution in which growth rate and transition probability are negatively correlated (lineages comprised of rapidly growing cells tend to transition between phases at a low rate (and vice versa)). During the simulation, lineages passed through the sequence of alternating Maturation and Dispersal Phases separated by sampling bottlenecks. During each phase, lineages grew exponentially until the total cell population of each reached carrying capacity. Additionally, each lineage may switch phenotype, with a probability defined by the corresponding trait value. Lineages that switch are established with the same parameters, but carry the new phenotype. Only lineages that contain a sub-lineage in which the phenotype has switched proceed to the next life cycle phase. Over time, some lineages go extinct due to competition and these are replaced with lineages from the same population. Hence, the distribution of traits across population changes with time. Evolution was recorded and analysed over 20 full cycles with 600 independent simulations.

The results show that cell growth rate (a proxy for cell fitness) slowly decreased in the Non-Mixed Ecology (Figure 7a), and rapidly increased in the Mixed Propagule Ecology (Figure 7b). At the same time, the transition probability (a proxy for group fitness) increased in the Non-Mixed Ecology (Figure 8a), while it remained stable in the Mixed Ecology (Figure 8b). Therefore, the model, comprising a minimal model in which evolution affects solely cell growth rate and capacity to switch phenotype, demonstrates that mixed and non-mixed regimes lead to qualitatively different evolutionary outcomes. Additionally, the simulations confirm that the pooling of propagules in the Mixed Propagule Ecology strengthens selection for the trait improving cell fitness (growth rate), which occurs at the expense of traits improving group fitness (transition probability).

**Figure 7.**
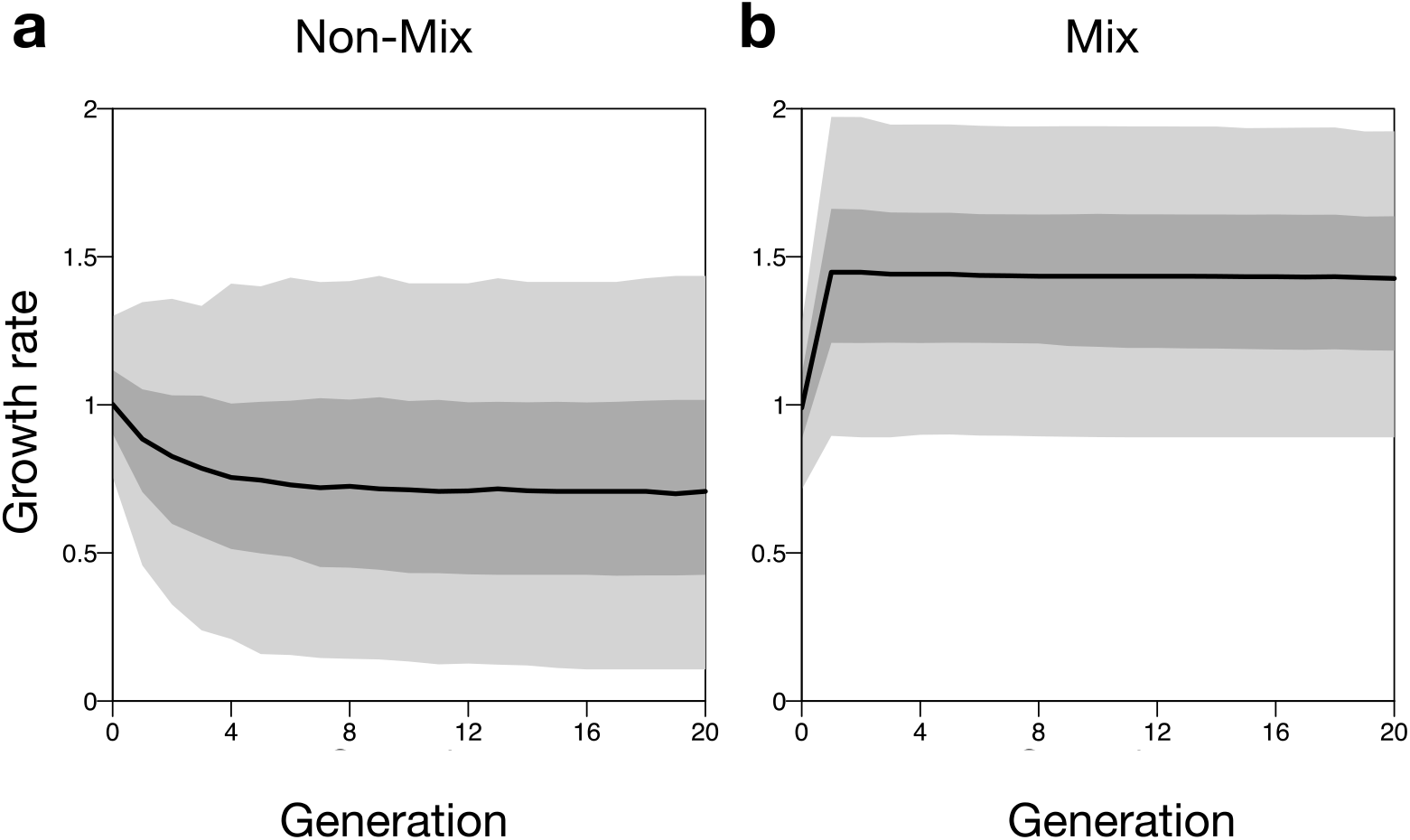
Simulated dynamics of the average cell growth rate in (a) the Non-Mixed Ecology, and (b) the Mixed Propagule Ecology. Black lines represent median growth rate values across 600 independent realizations of the respective selection regimes. Dark grey areas indicate a 50% confidence interval, while light grey areas indicate a 95% confidence interval.

**Figure 8.**
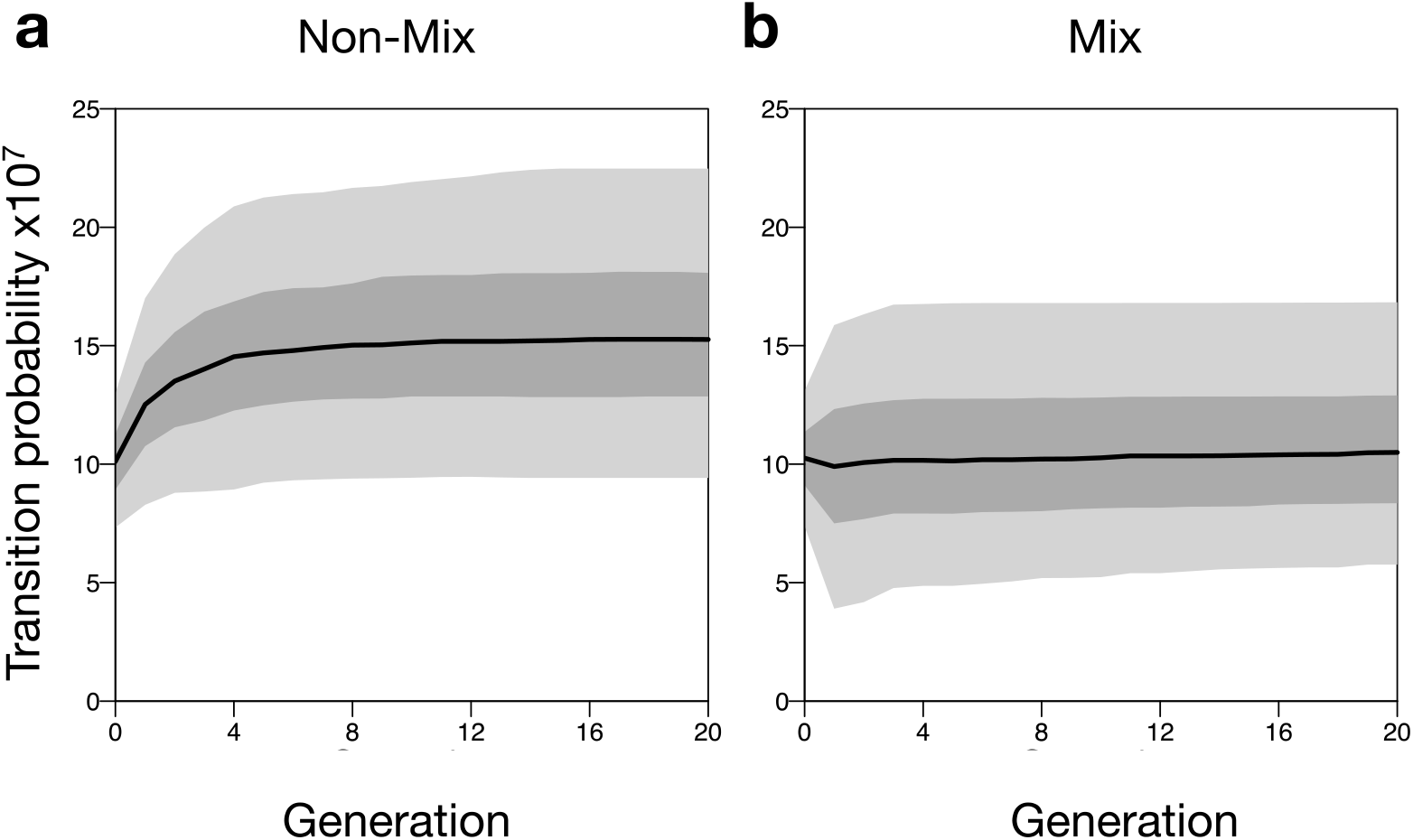
Simulated dynamics of the average transition probability in (a) the Non-Mixed Ecology, and (b) the Mixed Propagule Ecology. Black lines represent median transition rate values across 600 independent realizations of the respective selection regime. Dark grey areas indicate a 50% confidence interval, while light grey areas indicate a 95% confidence interval.

Given formulation of the model we asked whether eliminating the tradeoff between growth rate and transition probability affected the response of the evolving lineages to selection. Under the Non-Mixed Ecology, cell growth rate remained essentially unaffected (Supplementary Figure 1a), whereas with the tradeoff, cell growth rate declined (Figure 8a). In the Mixed Propagule Ecology, cell growth – in the absence of the tradeoff – remained as seen with the tradeoff (cf. Figure 8b with Supplementary Figure 1b). Under both non-mixed and mixed regimes transition probability (group fitness) increased although the increase in the mixed regime was smaller (Supplementary Figures 1c and 1d)). Together, results of the simulations are in full agreement with the experimental findings and emphasise the importance of the tradeoff between transition rate and WS density as evident in the experimental data.

### Summary of principle findings

Table 1 shows differences between ecologies in the partitioning of variation across meta-populations, including downstream consequences, for traits under selection in the non-mixed and mixed regimes. Selection during both phases of the Non-Mixed Ecology favoured a higher WS-SM transition rate. However, under the Mixed Propagule Ecology, the tradeoff between WS density and transition rate evident in the ancestral genotype, limited ability of selection to work on the collective lifecycle. Rather than acting on the life cycle as a whole, selection disproportionately affected cell-level selection. Adaptations may arise that allow groups to survive the Maturation Phase of the Mixed Propagule Ecology *(i.e.,* high WS-SM transition rate), only to be extinguished during the Dispersal Phase, due to a low competitive ability resulting from reduced WS density. The red box highlights the conflict between the effects of selection on the two incompatible traits involved in the two phases of the life cycle. This is further illustrated in Figure 9.

**Figure 9.**
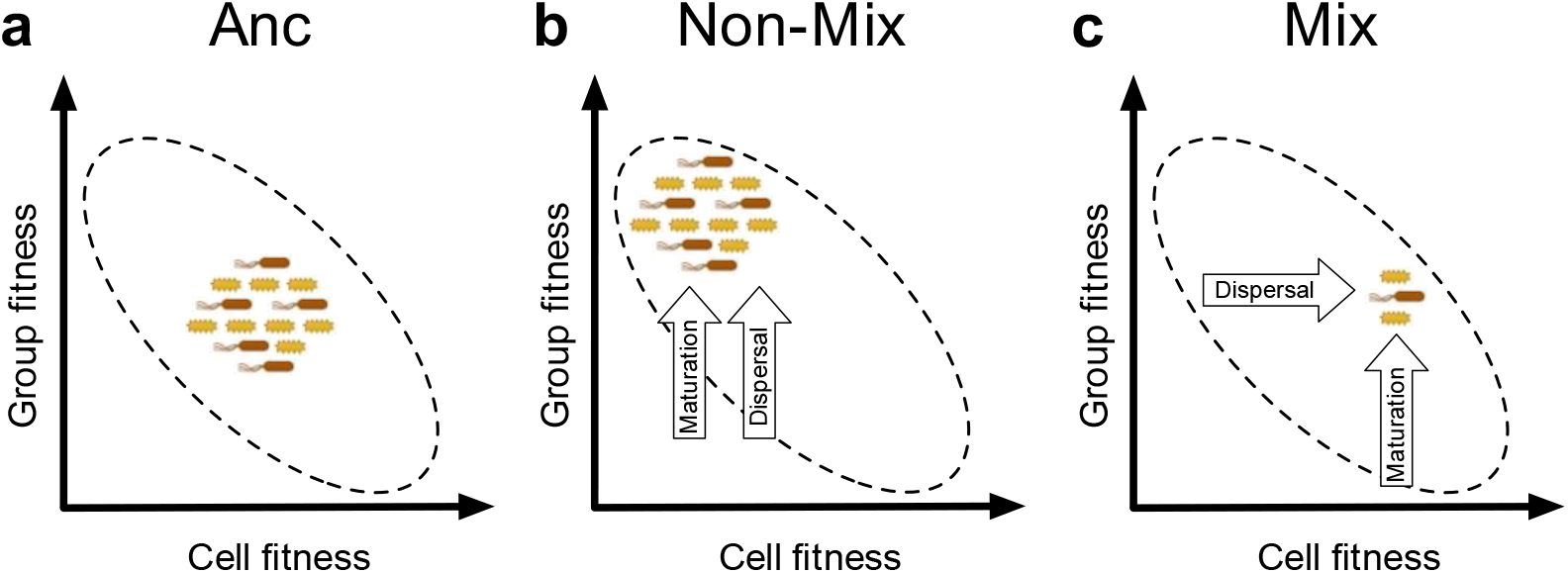
Ecological conditions can steer the evolution of traits in opposite directions. A metapopulation is depicted by the cloud of cells, where the position of the cloud represents the average fitnesses and the size of the cloud represents the diversity within a meta-population. The group and cell fitness are subjected to a tradeoff (dashed line eclipses) and cannot be optimized simultaneously. Arrows indicate the direction of selection applied by Maturation and Dispersal Phases of the life cycle. Both phases selected for increased transition rate in the Non-Mixed Ecology, whereas in the Mixed Propagule Ecology the Dispersal Phase promoted increased cell numbers at the expense of transition rate. The mixing procedure resulted in significantly decreased diversity, which further limited opportunity for adaptive evolution of groups.

**Table 1.**
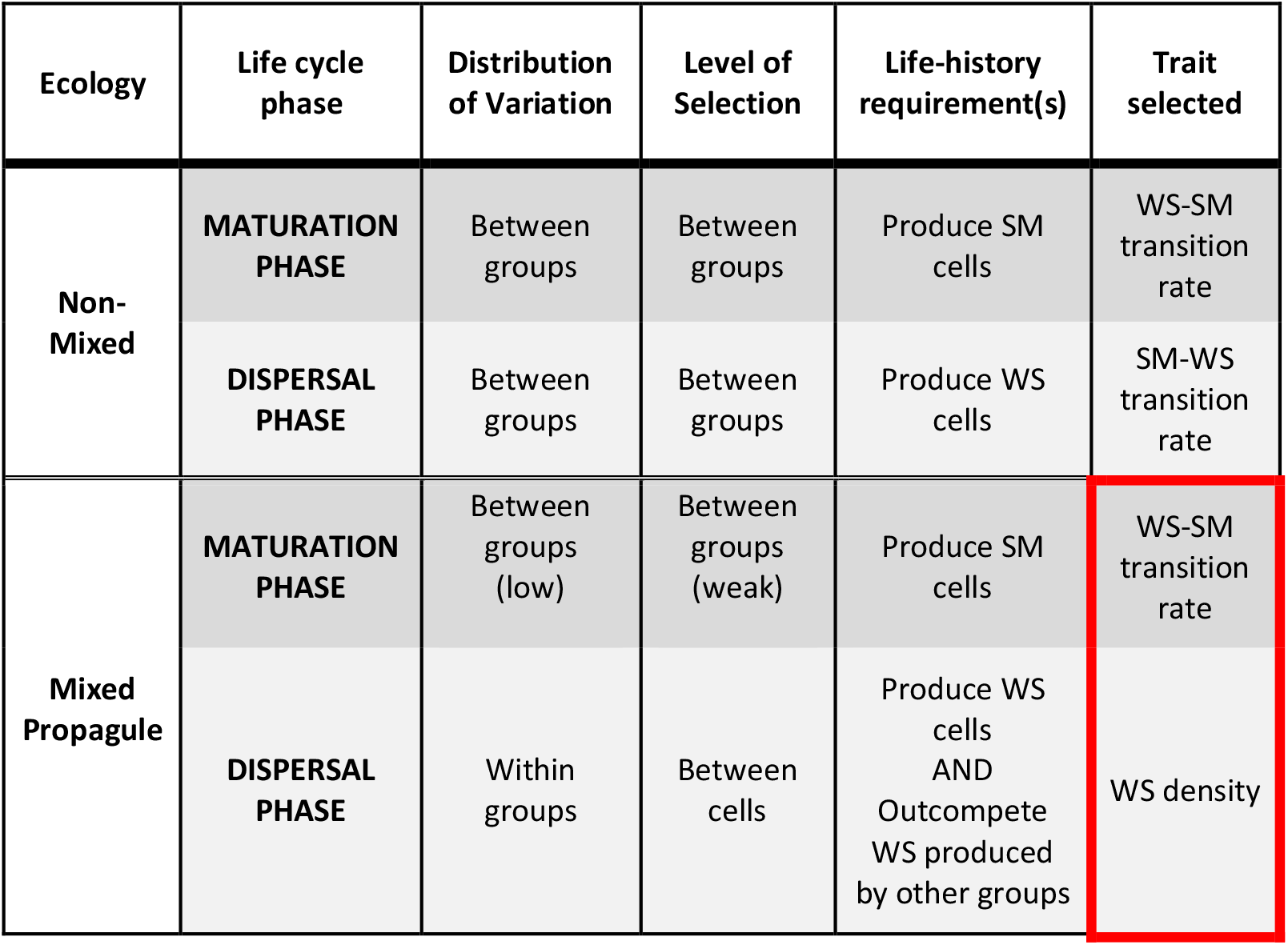
Effects of the meta-population structure on the level of selection. The red box highlights selection during different phases of the Mixed Propagule Ecology for two incompatible traits (parameters that are negatively correlated), leading to a conflict between levels of selection.

| Ecology | Life cycle phase | Distribution of Variation | Level of Selection | Life-history requirement(s) | Trait selected |
| --- | --- | --- | --- | --- | --- |
| Non-Mixed | MATURATION PHASE | Between groups | Between groups | Produce SM cells | WS-SM transition rate |
|  | DISPERSAL PHASE | Between groups | Between groups | Produce WS cells | SM-WS transition rate |
| Mixed Propagule | MATURATION PHASE | Between groups (low) | Between groups (weak) | Produce SM cells | WS-SM transition rate |
|  | DISPERSAL PHASE | Within groups | Between cells | Produce WS cells AND Outcompete WS produced by other groups | WS density |

### Perspective

Life cycles underpin evolutionary transitions to multicellularity (Buss 1987; Rainey 2007; Rainey and Kerr 2010; Hammerschmidt et al. 2014)). Life cycles solve the problem of group-level reproduction and shaped organismal form (Figure 10) (Buss 1987; Godfrey-Smith 2009; Rainey and Kerr 2010; Libby and Rainey 2013a; van Gestel and Tarnita 2017). Furthermore, life cycles involving reproductive specialisation provide selection with opportunity to act on something altogether novel – a developmental programme – that likely underpinned the rise of complexity in plants, animals and fungi (Grosberg and Strathmann 2007). Of further and particular significance is that life cycles establish the possibility that selection operates over a timescale longer than that of the doubling time of cells (Black et al. 2020). When this is accompanied by a death-birth process over the timescale of the life cycle, then selection over this longer timescale trumps within life cycle selection resulting in the fitness of groups decoupling from fitness of the composite cells. In the long-term, successful groups are composed of cells whose reproductive fate aligns with that of the longer time scale. This is the essence of the evolutionary transition from cells to multicellular life.

**Figure 10.**
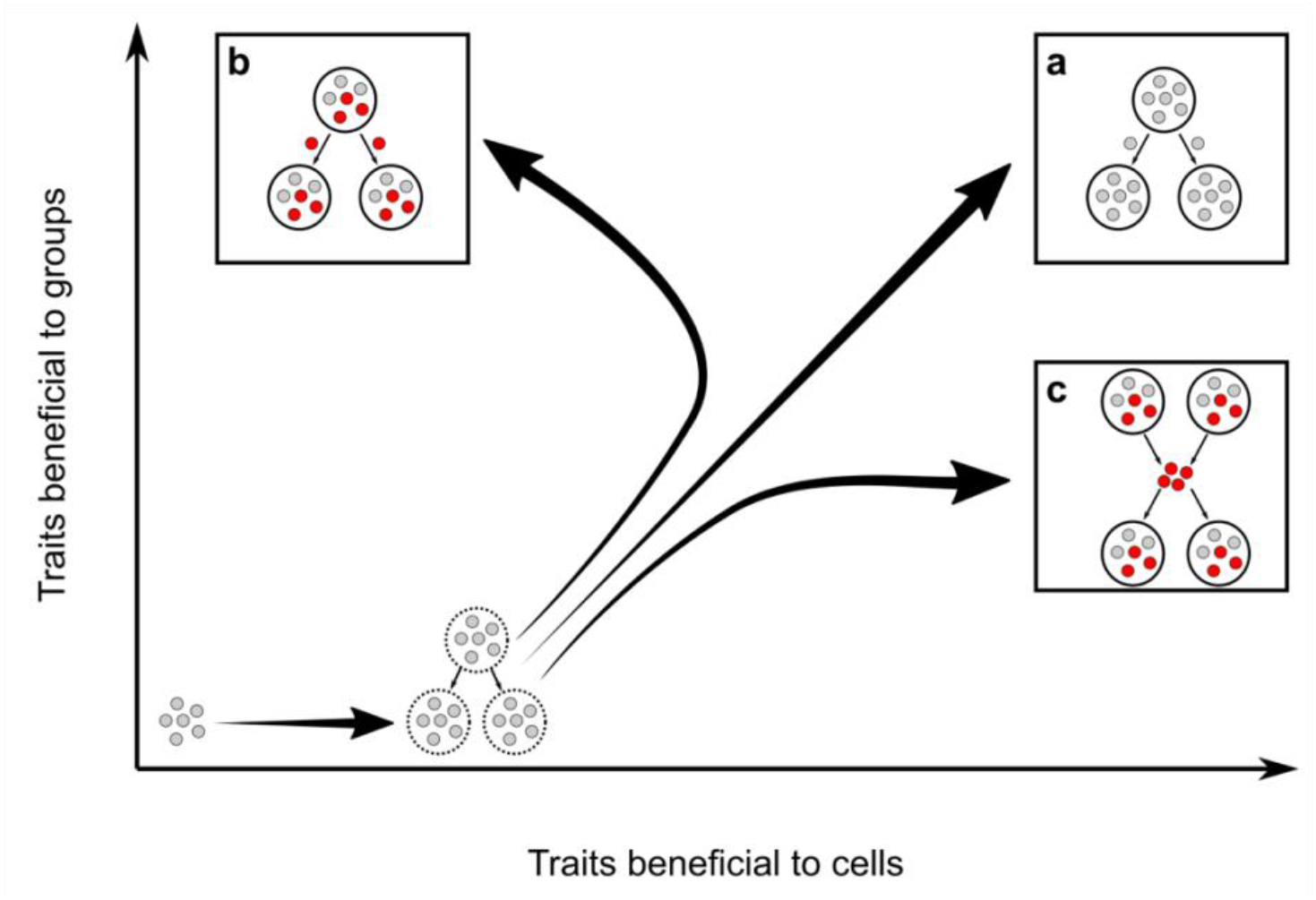
The origins of life cycles and the notion of fitness decoupling. Mode of group reproduction via a) fragmentation, b) a germ line (red) in a highly structured population and c) a germ line with propagule mixing, affects the emergence of individuality. Mode of group reproduction impacts the relationship between two levels of selection: the cell level (relative to the free-living state), and that of the emerging group. a) illustrates an example of a group that reproduces by fragmentation where fitness is ‘coupled’: group fitness is a by-product of the fitness of the constituent cells. Larger groups contain more cells and produce more offspring. This holds even when the reproductive life cycle involves a single-celled bottleneck – a feature that is expected to reduce within-group competition. b) and c) show examples of groups that reproduce via a life cycle involving two cell types – one soma-like and the other germ-like. Such two-phase life cycles allow possibility for traits determining a necessary developmental programme to evolve independent of the growth rate of cells that comprise the nascent organism. This paves the way for the emergence of new kinds of biological individual where group fitness ‘decouples’ from cell fitness.

It is instructive to place the findings from this study in the context of different modes of group reproduction and consequences for the expected long-term relationship between cell and group fitness. Figure 10 contrasts reproduction of groups via protected (unmixed) propagule lineages, unprotected (mixed) propagule lineages and fragmentation. In the latter, group fitness and cell fitness remain aligned. When propagules never mix, selection at the group level overwhelms cell-level selection, whereas when propagules mix, selection at the cell-level selection is the predominate driver of future evolutionary change.

When viewed among the diverse manifestations of multicellular life, non-mixing of propagules appears to be important for groups to begin the evolutionary trajectory toward paradigmatic forms of multicellularity, such as seen in metazoans. When propagules mix, our findings suggest the route toward less integrated forms of multicellularity as seen, for example, in the social amoeba, is more likely. Fragmentation of groups by equal division is problematic, resulting in the formation of chimeric organisms rife with cell-level conflict, and tellingly is exceedingly rare among multicellular life and found, to our knowledge, in Trichoplax alone.

True slime molds (Myxomycetes), and social Myxobacteria exhibit sophisticated behaviours such as ‘wolf-pack feeding’ that allow cells to benefit from group-living (Bonner 1998). Cellular slime molds such as the dictyostelids can form multicellular fruiting bodies when their food supply is exhausted (Strmecki et al. 2005). Organisms such as these exhibit rudimentary multicellular life cycles with cellular differentiation, and yet they have remained relatively simple for millions of years. This may be due, at least in part, to ecological factors that maintain a high degree of competition between cells from different groups during the single-cell phases of their respective life cycles. It is also likely that the aggregative mode of group formation (“coming together”) inhibits the process of selection at the aggregate level, compared to groups that form by growth from propagules (“staying together”) (Tarnita et al. 2013). It is interesting to note that in the experiments presented here, the benefits (to group fitness) of staying together were negated in the Mixed Propagule Ecology, which bear more resemblance to the coming together mode of group organisation during the Dispersal Phase of the life cycle.

If non-mixing among propagules is important for selection to work with potency on groups, then attention turns to environments and ecological circumstances that might ensure discreteness of the reproductive phase. Conceivably certain kinds of structured environments, such as found within soil pores might suffice. An alternate set of possibilities exist in environments where the density of propagules is low. For example, in the pond-plus-reed example that inspired our experimental studies, low nutrient levels in the pond may be sufficient to limit between-propagule competition.

Given a period of selection for traits that favour the persistence of groups, more integrated collectives may withstand a less structured ecology. In other words, a structured environment can provide the ecological scaffold necessary to support persistence during an initial period of evolution in which complex adaptations arise and prevail over selection solely for growth rate. Upon removal of the scaffold, features such as boundaries that demarcate groups, would allow collectives to continue to function as evolutionary individuals (Black et al. 2020).

Extant multicellular organisms tolerate varying degrees of cell-level selection, as evidenced by the diverse modes of multicellular reproduction that incorporate intense competition at the gamete level. Many plants, for example, engage in synchronous seed dispersal – a life cycle not unlike that depicted in Figure 10c. Cancer is a classic example of lower-level selection occurring in many multicellular organisms that is largely contained by selection at the higher level (cancers generally arise later in life, after reproduction (Nunney 1999)). In polyandrous animals, sexual selection also occurs at two levels: a higher level with competition between individuals for mating, and a lower level with competition between sperm for fertilization of eggs within female genital tracts. This lower level has often been shown to account for a large fraction of total variance in male fitness (and hence of the opportunity for selection); for example, 46% in red jungle fowl (Collet et al. 2012), or 40% in snails (Pélissié et al. 2014). Competition between units of the lower level (i.e., germ cells) is extreme in many aquatic invertebrates during broadcast spawning. Here, the animals (higher level) never meet, as sperm and eggs (lower level) are released into the water column, where competition among the gametes for fertilization takes place.

Given the unknown evolutionary history of organisms that reproduce by life cycles in which there is intense cell-level selection, and the seeming incompatibility of such modes of reproduction with our experimental findings, it is important to recognise that such modes are likely derived and determined by ecological conditions experienced after nascent multicellular forms arose. This draws attention to a possible alternate solution for minimising propagule-level competition that stems from development.

Assuming discreteness of the group phase, and opportunities for group-level dispersal, then collectives that evolve capacity to retain the propagule phase as an integral part of the group, releasing newly created offspring only after the multicellular (albeit immature) state has been achieved, would likely fare well. Such groups would experience minimal between-group selection at the single cell stage and selection would be predominantly group-level. That this mode of reproduction is a feature of paradigmatic forms of multicellularity – along with the fact that germ cells do not typically replicate once produced – likely marks the importance of early developmental innovations for the evolution of complex multicellular life.

## Data accessibility

All data are available at https://zenodo.org/record/3748416#.XpGdTC17Fgg.

## Supplementary material

A single supplementary figure is appended below

## Acknowledgements

We thank B. Kerr and E. Libby for discussion, P. Bourrat for comments on drafts of the manuscript and the sterling efforts of D. Brisson in his editorial role. The work was directly supported by the Marsden Fund Council from government funding administered by the Royal Society of New Zealand, and in part by grant RFP-12-20. Version 5 of this preprint has been peer-reviewed and recommended by *PCI Evol Biol* (https://doi.org/10.24072/pci.evolbiol.100099).

## Author contributions

K.H., C.J.R, and P.B.R. contributed to the conception and design of the study. K.H. and C.J.R. performed research, undertook data analysis and prepared figures. Y.P. designed the numerical model, performed simulations and prepared figures. All authors wrote the paper.

## Supplementary figure

**Supplementary Figure 1.**
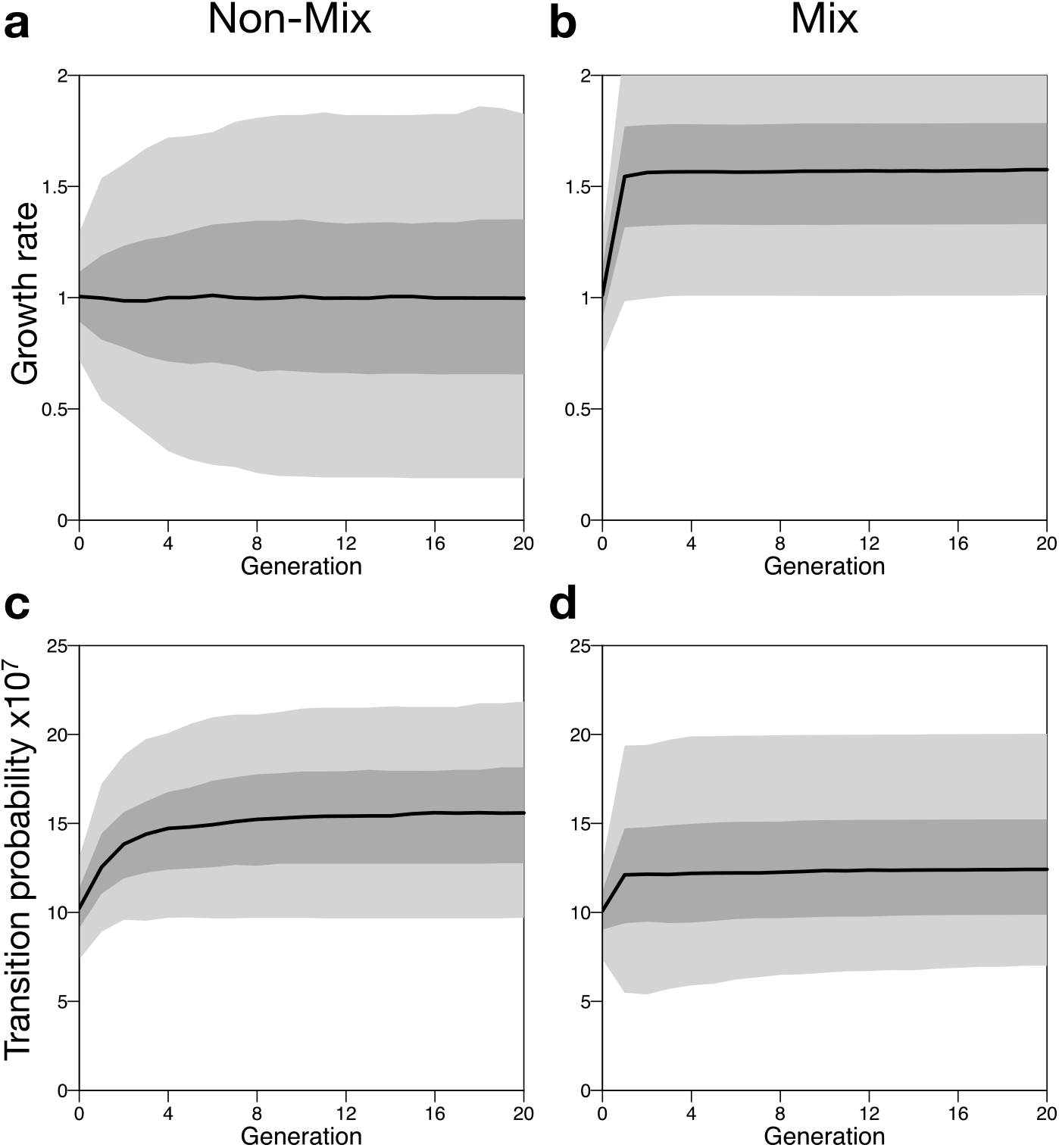
Model dynamics with a random distribution of initial parameters (no tradeoff) (a,b) average cell growth rate, and (c,d) average transition probability in the Non-Mixed Ecology (a,c) and the Mixed Propagule Ecology (b,d). Black lines represent median values across 600 independent realizations of the respective selection regime. Dark grey areas indicate a 50% confidence interval, while light grey areas indicate a 95% confidence interval.

## References

Black AJ, Bourrat P, Rainey PB. 2020. Ecological scaffolding and the evolution of individuality. Nat Ecol Evol. 4:426–436.

Bonner JT. 1998. The origins of multicellularity. Integr Biol. 1:27–36.

Boraas ME, Seale DB, Boxhorn JE. 1998. Phagotrophy by a flagellate selects for colonial prey: a possible origin of multicellularity. Evol Ecol. 12:153–164.

Bourke AF. 2011. Principles of Social Evolution. Oxford: Oxford University Press

Bourrat P. (2015) Levels, time and fitness in evolutionary transitions in individuality. Phil Theor Biol. 7 doi:10.3998/ptb.6959004.0007.001

Buss LW. 1987. The Evolution of Individuality. Princeton University Press

Collet J, Richardson DS, Worley K, Pizzari T. 2012. Sexual selection and the differential effect of polyandry. Proc Natl Acad Sci USA 109:8641–8645.

De Monte S, Rainey PB. 2014. Nascent multicellular life and the emergence of individuality. J Biosci. 39:237–248.

Godfrey-Smith P. 2009. Darwinian Populations and Natural Selection. Oxford University Press

Griesemer J. 2001. The units of evolutionary transition. Selection 1:67–80.

Grosberg RK, Strathmann RR. 2007. The evolution of multicellularity: A minor major transition? Annu Rev Ecol Evol Syst. 38:621–654.

Hammerschmidt K, Rose CJ, Kerr B, Rainey PB. 2014. Life cycles, fitness decoupling and the evolution of multicellularity. Nature 515:75–79.

Herron MD, Borin JM, Boswell JC, Walker J, Chen I-CK, Knox CA, Boyd M, Rosenzweig F, Ratcliff WC. 2019. De novo origins of multicellularity in response to predation. Sci Rep 9:2328.

Libby E, Rainey PB. 2013a. A conceptual framework for the evolutionary origins of multicellularity. Phys Biol. 10:035001.

Libby E, Rainey PB. 2013b. Eco-evolutionary feed-back and the tuning of proto-developmental life cycles. PLoS One, DOI: 10.1371/journal.pone.0082274.

Lind PA, Farr AD, Rainey PB. 2015. Experimental evolution reveals hidden diversity in evolutionary pathways. eLife 4:441.

Lind PA, Farr AD, Rainey PB. 2017. Convergence in experimental *Pseudomonas* populations. ISME J 11:589–600.

Lind PA, Libby E, Herzog J, Rainey PB. 2019. Predicting mutational routes to new adaptive phenotypes. eLife 8:e38822.

McDonald MJ, Gehrig SM, Meintjes PL, Zhang XX, Rainey PB. 2009. Adaptive divergence in experimental populations of *Pseudomonas fluorescens*. IV. Genetic constraints guide evolutionary trajectories in a parallel adaptive radiation. Genetics 183:1041–1053.

Michod RE, Roze D. 1999. Cooperation and conflict in the evolution of individuality. III. Transitions in the unit of fitness. In: Mathematical and Computational Biology Computational Morphogenesis, Hierarchical Complexity, and Digital Evolution. (ed. Nehaniv CL) pp. 47–92.

Nunney L. 1999. Lineage selection and the evolution of multistage carcinogenesis. Proc R Soc B. 266:493–498.

Pélissié B, Jarne P, Sarda V, David P. 2014. Disentangling precopulatory and postcopulatory sexual selection in polyandrous species. Evolution 68:1320–1331.

Queller DC, Strassmann JE. 2009. Beyond society: the evolution of organismality. Phil Trans R Soc B. 364:3143–3155.

Rainey PB, De Monte S. 2014. Resolving conflicts during the evolutionary transition to multicellular life. Annu Rev Ecol Evol Syst. 45:599–620.

Rainey PB, Kerr B. 2010. Cheats as first propagules: A new hypothesis for the evolution of individuality during the transition from single cells to multicellularity. BioEssays 32:872–880.

Rainey PB, Rainey K. 2003. Evolution of cooperation and conflict in experimental bacterial populations. Nature 425:72–74.

Rainey PB, Travisano M. 1998. Adaptive radiation in a heterogenous environment. Nature 394:69–72.

Rainey PB, Remigi P, Farr AD, Lind PA. 2017. Darwin was right: where now for experimental evolution? Curr Opin Genet Dev. 47:102–109.

Rainey PB. 2007. Unity from conflict. Nature 446:616.

Rose CJ, Hammerschmidt K, Rainey PB. (2020). Analysis of an experimental transition in individuality challenges the need to assign traits to levels. bioRxiv: https://doi.org/10.1101/2020.03.02.973792

Silby MW, Cerdeño-Tárraga AM, Vernikos GS, Giddens SR, Jackson RW, Preston GM, Zhang X-X, Moon CD, Gehrig SM, Godfrey SA, et al. 2009. Genomic and genetic analyses of diversity and plant interactions of *Pseudomonas fluorescens*. Genome Biol. 10:R51.

Spiers AJ, Kahn SG, Bohannon J, Travisano M, Rainey PB. 2002. Adaptive divergence in experimental populations of *Pseudomonas fluorescens*. 1. Genetic and phenotypic bases of wrinkly spreader fitness. Genetics 161:33–46

Spiers AJ, Bohannon J, Gehrig SM, Rainey PB. 2003. Biofilm formation at the air-liquid interface by the *Pseudomonas fluorescens* SBW25 wrinkly spreader requires an acetylated form of cellulose. Mol Microbiol. 50:15–27

Strmecki L, Greene DM, Pears CJ. 2005. Developmental decisions in *Dictyostelium discoideum*. Dev Biol. 284:25–36.

Tarnita CE, Taubes CH, Nowak MA. 2013. Evolutionary construction by staying together and coming together. J Theor Biol. 320:10–22.

van Gestel J, Tarnita CE. 2017. On the origin of biological construction, with a focus on multicellularity. Proc Natl Acad Sci USA. 114:11018–11026.

West SA, Fisher RM, Gardner A, Kiers ET. 2015. Major evolutionary transitions in individuality. Proc Natl Acad Sci USA. 112:10112–10119.

West SA, Griffin AS, Gardner A. 2007. Evolutionary explanations for cooperation. Curr Biol. 17:R661–R672.

Wilson DS. 1975. A theory of group selection. Proc Natl Acad Sci USA. 72:143–146.

Zhang X-X, Rainey PB. 2007. Construction and validation of a neutrally-marked strain of *Pseudomonas fluorescens* SBW25. J Microbiol Meth. 71:78–81.

